# Oncogenic lncRNA transgene transcription modulates epigenetic memory at a naïve chromosomal locus

**DOI:** 10.1101/2025.05.15.654293

**Authors:** Sweta Sikder, Songjoon Baek, Yamini Dalal, Ganesan Arunkumar

## Abstract

Maintaining genome integrity is crucial for the proper functioning and development of organisms. One intriguing aspect of genome integrity is the formation and function of neocentromeres at non-centromeric sites. CENP-A, a centromere-specific protein, is essential for centromere identification and function. However, in many cancers, CENP-A is often found to be ectopically misplaced when overexpressed. Moreover, CENP-A deposition at the centromere depends on the transcription of centromeric non-coding RNAs. Consequently, ectopic CENP-A is found at transcriptionally active and frequent breakpoint regions. To further explore ectopic CENP-A localization, we previously engineered a stable ectopic CENP-A site on a naïve chromosome by overexpressing a non-centromeric oncogenic lncRNA, PCAT2, which was capable of recruiting CENP-A to its transcription site. In this work, we tracked cells carrying this stable transgene to understand the longevity of the induced ectopic CENP-A site at the chromosome that harbors it. Our findings revealed that the induced epigenetic memory was eventually lost due to the suppression of the transgene through competing epigenetic silencing mechanisms. This epigenetic restoration naturally reversed the ectopic CENP-A level to its previous levels at the engineered site. These data suggest that cells may have evolved failsafe mechanisms to prevent neocentromere formation at ectopic sites by suppressing transcription, unless otherwise favored by selection involving multiple components.

## Introduction

Genome integrity is crucial for the proper functioning and development of an organism. Any disruptions or alterations in genome integrity can lead to genetic disorders or diseases. One intriguing aspect of genome integrity is the formation of neocentromeres. Centromeres are specialized regions on chromosomes responsible for ensuring accurate chromosome segregation during cell division. Neocentromeres, conversely, are newly formed centromeres that arise at non- canonical genomic loci, typically devoid of centromeric DNA sequences^1^. The formation of neocentromeres often occurs in response to genetic alterations, such as chromosomal rearrangements, deletions, or translocations^2^. However, the mechanisms underlying the establishment and maintenance of neocentromeres remain an exciting area of exploration. Epigenetic modifications, such as DNA methylation, histone modifications, and histone variant mislocalization, are intricately involved in neocentromere formation^3–5^.

CENP-A is a histone H3 variant which epigenetically marks centromeres, ensures faithful chromosome segregation, and maintains genome integrity through cell divisions^6^. In mammals, the histone chaperone HJURP facilitates the specific incorporation of CENP-A into centromeric chromatin during late mitosis in a transcription-dependent pathway independent of S phase^7,8^. CENP-A consistently localizes to the same centromeric sequences before and after DNA replication^9^. Following the deposition of the CENP-A nucleosome at the centromeric DNA sequence, CENP-A utilizes its C-terminal tail to directly bind CENP-C in two domains within CENP-C^10,11^. CENP-C binding to CENP-A stabilizes centromeric chromatin, rigidifies both surface and internal nucleosome structure, and unwraps the terminal DNA of CENP-A^10,12,13^. Intriguingly, in several multicellular species, CENP-A alone can drive centromeric identity independent of a defined centromeric DNA sequence^14,15^. Therefore, deposition of CENP-A is carefully regulated to occur only once per cell cycle at the centromeres, taking place during the early G1 phase^7,8,16,17^. Deposition of CENP-A into centromeric chromatin was spatiotemporally restricted through various mechanisms, including the recruitment of licensing factor Mis18BP1 to centromeres early during mitosis, depending on Plk1-mediated phosphorylation^18–20^. To prevent erroneous deposition of CENP-A to non-centromeric regions, CENP-A protein levels are tightly controlled by specific E3 ubiquitin ligase-mediated proteolysis^21,22^. CENP-A is detectable at low levels in highly transcribed euchromatic regions of the yeast genome and on overexpression localized to euchromatic regions in certain organisms^9^. In addition, CENP-A can also be detected at the sites of DNA double-strand breaks before being removed during DNA repair^23^. However, these findings suggest that although CENP-A can sporadically localize to euchromatin, it is actively maintained only at centromeres^9,21^. Nevertheless, CENP-A overexpression has been identified in many cancers, including breast, central nervous system, colorectal, lung, and ovary^3,6^. This overexpression can have various effects on the cell, including changes in gene expression, genome integrity, reprogramming of cell fate, and alterations in 3D nuclear organization^6,24–27^. Importantly, high levels of CENP-A lead to its mislocalization outside of the centromeres^26–31^. CENP-A mislocalization to a non-centromeric site could lead to the formation of neocentromeres, which could compete with the original centromere function^29,32–34^. These ectopic CENP-A domains are found to be colocalized with binding partners, such as CENP-C^29,30,32,35^, suggesting that they can form kinetochores^29^. This, in turn, can contribute to chromosome instability^29,30,32,36^.

CENP-A loading to centromeres require active transcription of centromeric alpha-satellite DNA in many species^37^. RNA pol II transcription and centromere-derived long non-coding RNAs (lncRNA) physically interact with CENP-A and its chaperone HJURP to deposit CENP-A to the centromeric DNA^38–42^. LncRNAs represent a class of transcribed RNA molecules that are longer than 200 nucleotides and with limited or no coding potential. LncRNA plays crucial roles in diverse physiological and pathological functions; therefore, it is precisely regulated by epigenetic mechanisms and other molecules^43^. Dysregulation of lncRNA expression correlates with tumorigenesis, invasiveness, and drug resistance through diverse mechanisms in multiple types of cancer^44^. LncRNAs can interact with DNA, RNA, and proteins through base pairing or the functional domains that arise through their secondary and tertiary structures. As versatile molecules, they can form physical and functional interactions with these biomolecules through *cis* and *trans* mechanisms^43^. For instance, centromeric and pericentromeric non-coding RNAs interact with various centromeric and kinetochore components to establish an active and functional centromere-kinetochore network^37^. Interestingly, some non-centromeric lncRNAs also function in *trans* to organize and regulate centromeric function^45^. This raises the question: could non-centromeric lncRNAs function as recruiting competitor signals for centromeric proteins at ectopic sites?

Previously, we reported that the oncogenic lncRNA *PCAT2,* encoded from the gene desert surrounding the *cMYC* locus on chromosome 8, plays a role in recruiting CENP-A to the 8q24 locus in SW480 metastatic colon cancer cells^32^. Interestingly, *PCAT2* also recruited CENP-A to an exogenously expressed *PCAT2* cassette from a transgene locus on chromosome 4 in human colon cancer cells, which naturally overexpress CENP-A due to the loss of p53 regulation^32^. Similar to centromeric lncRNAs, *PCAT2* physically co-purifies with CENP-A and CENP-C *in vivo*, with enrichment of R-loops at the site of transcription^32^. We took advantage of this transgene *PCAT2* (Trans*PCAT2*) locus on chromosome 4q31 to elucidate the mechanism by which the new CENP-A ectopic sites are epigenetically established, maintained, or lost in the CENP-A overexpressing colon cancer cell line. We find that induced epigenetic memory resulting from mislocalized ectopic CENP-A is transient, without persistent transcription of the Trans*PCAT2* lncRNA. Furthermore, levels of ectopic CENP-A at the Trans*PCAT2* locus were diminished over a year, while the innate *PCAT2* locus at chromosome 8q24 retained the steady-state level of ectopic CENP-A. We report that Trans*PCAT2* knock-in cells may have triggered an epigenetic silencing mechanism that involve differential DNA methylation and heterochromatin formation to suppress the expression of the oncogenic lncRNA transgene. This is similar to the epigenetic silencing mechanism we previously observed at aging centromeres^46^. By utilizing these epigenetic silencing mechanisms, the cell prevents the accumulation of ectopic CENP-A at the transgene locus, thereby inhibiting the formation of neocentromeres and maintaining chromosome stability.

## Results

### Transcriptionally inactive lncRNA Trans*PCAT2* locus lost ectopic CENP-A

Our previous study demonstrated that the oncogenic lncRNA *PCAT2*, transcribed from a non-native locus on chromosome 4q31, can form an ectopic CENP-A domain in a cancer background with CENP-A overexpression (Figure 1A)^32^. This Trans*PCAT2* locus at around week 16 has ectopic CENP-A levels equal to the 8q24 locus levels. Here, we sought to investigate whether this epigenetic memory remained stable over time and could contribute to the formation of a neocentromere-like domain in cancer cells. Over a two-year period, we tracked SW480*^PCAT2^*^-KI^ colon cancer cells engineered to carry three copies of the *PCAT2* gene at the chromosome 4q31 locus. We performed whole-genome sequencing of the CRISPR knock-in stable cells after the cell line establishment^32^ . Subsequently, there was an initial four-fold increase in ectopic CENP-A levels at the Trans*PCAT2* locus until week 20^32^. CENP-A was indirectly stained using CENP-C, that binds only to CENP-A and present in all CENP-A ectopic sites^29,32,36^. After about week 22, the CENP-A levels at this locus gradually reduced to their native levels by week 40 and subsequently remained stable (Figure 1B). In addition, we noted that the transcript levels of Trans*PCAT2* were significantly higher at week 16 compared to weeks one, 90 and 110 (Figure 1C). Therefore, we performed the whole-genome sequencing to confirm the presence of the Trans*PCAT2* transgene sequence at the 4q31 site (Figure S1A). We also performed CENP-A ChIP-seq to determine the levels of CENP-A at the Trans*PCAT2* locus at 4q31 and confirmed the reduction of the CENP-A signal at this locus (Figure 1D). The reduction in Trans*PCAT2* transcript levels correlated with the decrease in ectopic CENP-A levels at the Trans*PCAT2* locus (Figure 1B, D, & E). The ectopic CENP-A levels at the 8q24 locus remained unchanged in this cell line (Figure 1B & E, Figure S1B). These observations, in line with our previous findings^32^, suggest that active transcription is required for the deposition and maintenance of CENP-A and that the level of ectopic CENP-A at the Trans*PCAT2* locus depends on the active transcription of the Trans*PCAT2* lncRNA gene^32,38^.

**Figure 1.**
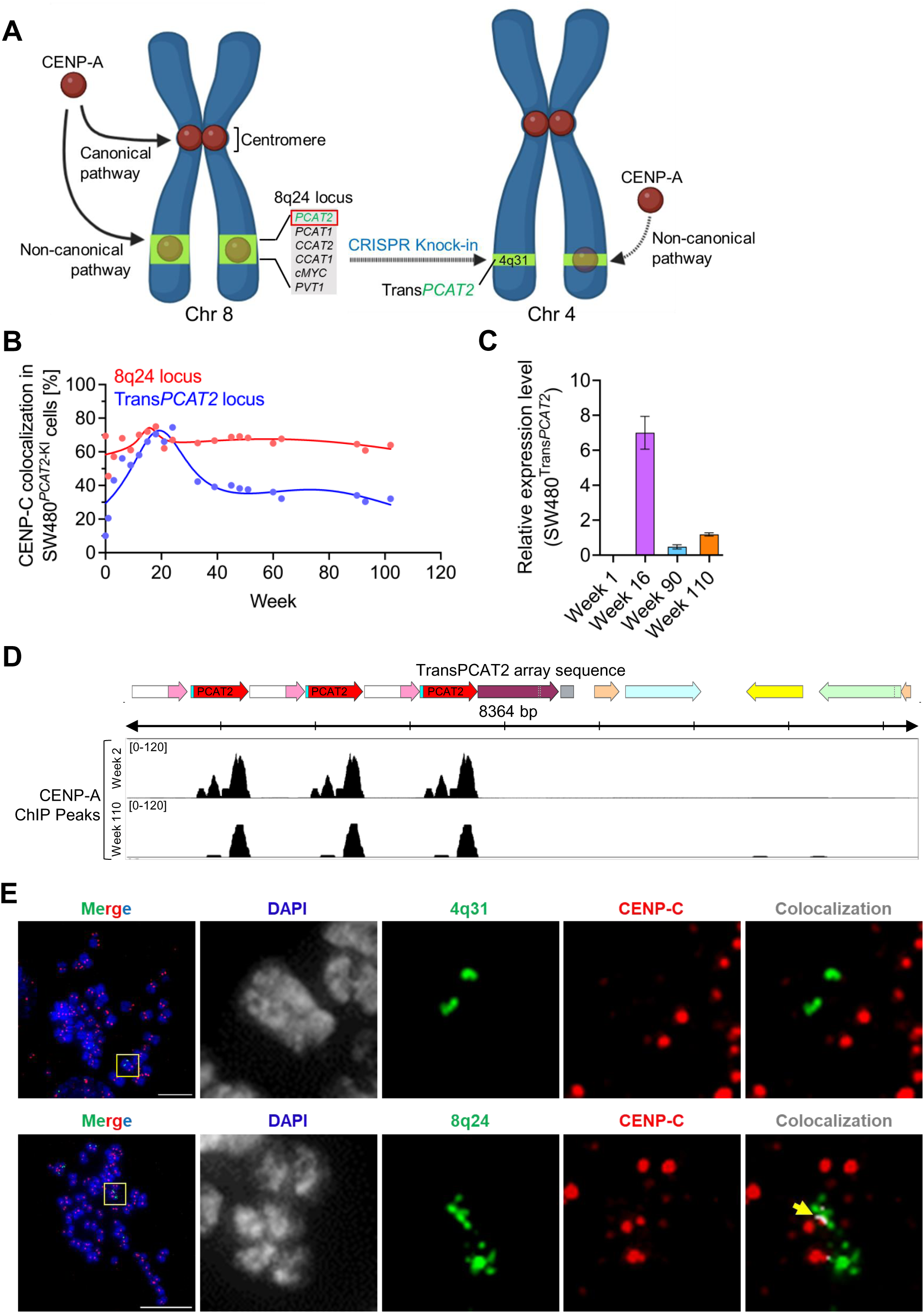
LncRNA *PCAT2*-mediated induced CENP-A ectopic site lost CENP-A over time. **(A)** Graphical illustration shows CENP-A ectopic deposition to a non-centromeric site, such as the 8q24 breakpoint, utilizing a non-canonical pathway where it forms a stable ectopic site. The 8q24 locus is enriched with several oncogenic lncRNAs involved in recruiting CENP-A. The transcription of 8q24-derived lncRNA *PCAT2* plays a significant role in recruiting CENP-A to its transcribing locus. Through CRISPR knock-in of a transgene cassette carrying *PCAT2* and its subsequent ectopic transcription from a naïve chromosome locus (4q31), CENP-A is recruited to the transgene inserted locus, ultimately forming a novel CENP-A ectopic site. **(B)** Line graphs showing the level of CENP-A at the 4q31 locus of SW480 wild and Trans*PCAT2* knock-in cells. CENP-A was stained indirectly using CENP-C, a binding partner of CENP-A in centromeres and ectopic CENP-A sites. The CENP-A levels peaked at week 20 and reduced gradually to their original levels after week 40. **(C)** Histogram showing the expression level of Trans*PCAT2* transcript in SW480*^PCAT2^*^-KI^ cells at different time points. **(D)** CENP-A ChIP peak browser track showing CENP-A occupancy at the TransPCAT2 array inserted at the 4q31 locus of SW480*^PCAT2^*^-KI^ cells displayed using a custom-built genome. The peak intensity demonstrates the decrease in CENP-A level at week110 compared to week 2 samples. **(E)** Immunofluorescence-DNA FISH image showing the loss of CENP-A signal from the Trans*PCAT2* gene site at 4q31 locus of SW480*^PCAT2^*^-KI^ cells at week 90. However, the 8q24 locus is still retaining CENP-A at this time point. CENP-A is indirectly stained with CENP-C (red), and the 4q31 locus is marked with locus-specific FISH probe (green). The scale bar represents 10 µm.

### Trans*PCAT2* insertion altered the transcription pattern of the 4q31 locus through rewiring the epigenetic landscape

We next tested if the insertion of the Trans*PCAT2* gene at the 4q31 locus altered the expression patterns of the adjacent genes. Therefore, we compared the expression of Trans*PCAT2* adjacent genes *MAML3*, *SCOC*, *SCOC-AS1*, and *SMAD1* at weeks 1, 16, and 90 (Figure 2A). Interestingly, compared to week 1, all tested 4q31 locus gene transcripts were undetectable at week 16 when the Trans*PCAT2* level was significantly higher, except for the anti- sense RNA *SCOC-AS1* (Figures 1C & 2B). However, by week 90, all 4q31 locus genes had returned to their native expression levels (Figure 2B). Notably, the genes *PLK4* in 4q28.1 and *DHX15* in 4p15.2, distant from the Trans*PCAT2* gene, also showed similar expression pattern change to the 4q31 locus genes (Figure 2B), highlighting that there was gene expression change in chromosome-wide. Subsequently, we performed total RNA-seq from week 110 SW480*^PCAT2^*^-KI^ cells and identified a significant change in the transcriptome. There are 13,276 genes differentially expressed (7,412 genes differentially expressed in week 2 and 5,864 in W110 with *p*<0.05) between SW480*^PCAT2^*^-KI^ cells from week 2 and week 110 (Figure S2). These findings prompt consideration of whether introducing an oncogenic lncRNA gene to a non-native site on a naïve chromosome fundamentally alters the expression of many genes in that chromosome.

**Figure 2.**
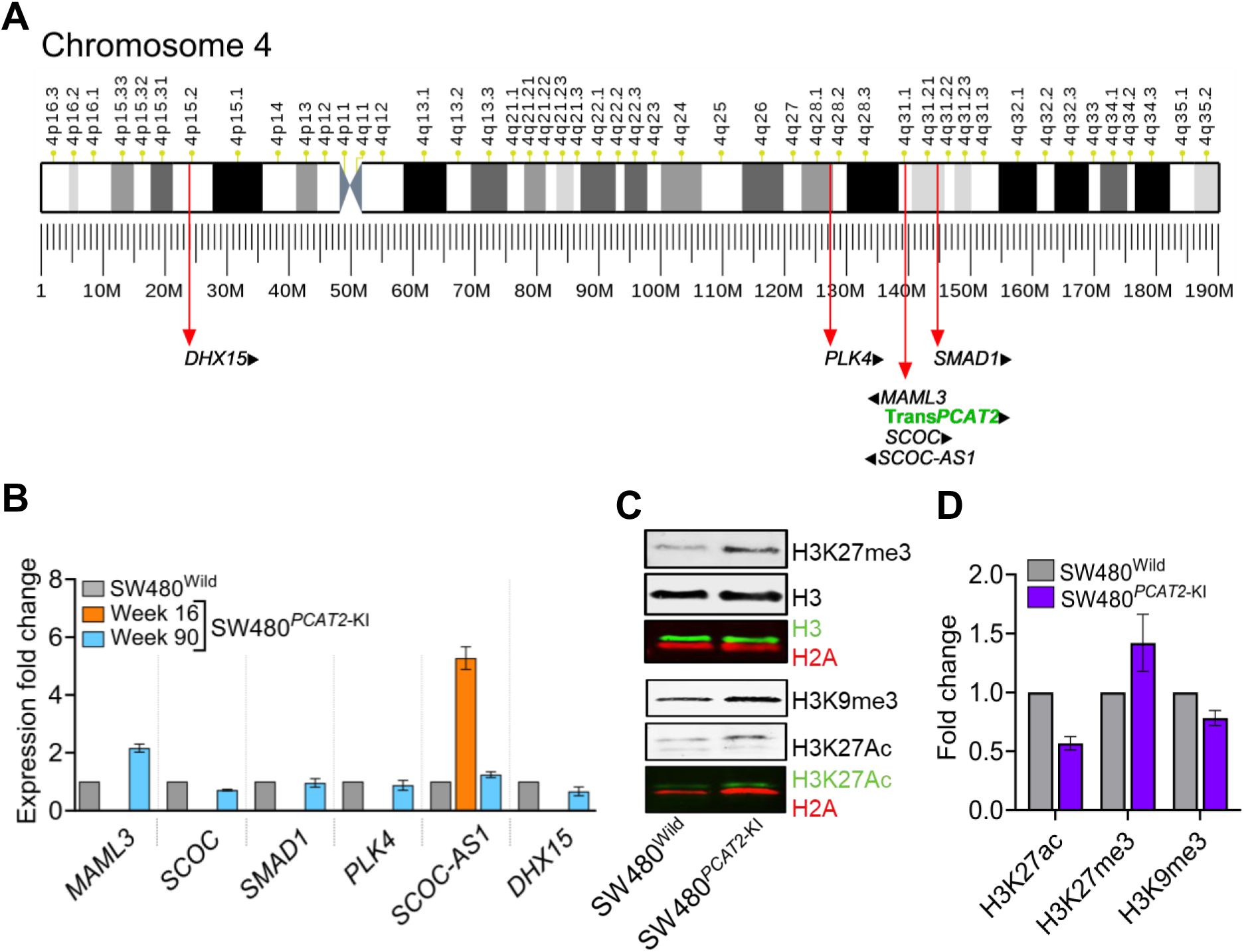
Introduction of *PCAT2* to non-native sites and its expression impacted global epigenetic signature. **(A)** The ideogram depicts the knock-in position of the Trans*PCAT2* gene into the chromosome 4q31 locus of SW480 cells, with reference to other control genes studied in this work. **(B)** Histogram displays the expression levels of the Trans*PCAT2* transcript and the proximal (*MAML3*, *SCOC*, and *SCOC-AS1*) and distal genes (*SMAD1*, *PLK4*, and *DHX15*) relative to the knock-in site of the Trans*PCAT2* gene at week 16 and 90 in SW480*^PCAT2^*^-KI^ cells, in comparison to SW480^Wild^ cells. **(C)** Representative images of immunoblots show the levels of H3K27me3, H3K9me3, and H3K27ac marks in SW480*^PCAT2^*^-KI^ cells in comparison to SW480^Wild^ cells. **(D)** Histogram shows the quantification of various epigenetic marks between SW480^Wild^ and SW480*^PCAT2^*^-KI^ cells shown in the immunoblots.

Epigenetic silencing, facilitated by the formation of heterochromatin, has been linked to gene regulation and chromosome integrity. Studies have shown that introducing transgenes near heterochromatin regions can exhibit a variegating phenotype^47^. To test whether the Trans*PCAT2* bearing 4q31 locus is heterochromatinized, we isolated nuclei from SW480^Wild^ and SW480*^PCAT2^*^-KI^ cells and examined the global epigenetic regulatory markers H3K9me3 and H3K27me3, and activation mark H3K27ac (Figure 2C). Interestingly, all the epigenetic marks tested were dysregulated in the SW480*^PCAT2^*^-KI^ cells compared to SW480^Wild^ cells (Figure 2C-D). We observed that H3K27ac and H3K9me3 levels decreased modestly, while the H3K27me3 mark was significantly increased genome-wide in the SW480*^PCAT2^*^-KI^ cells compared to the wild cells (Figure 2C-D). Our results demonstrate that knocking-in and constitutive overexpression of oncogenic lncRNA Trans*PCAT2* at a non-native chromosome site could significantly affect the global epigenetics landscape and induce heterochromatin reorganization.

To understand the global epigenetic changes, we performed ChIP-seq between SW480*^PCAT2^*^-KI^ and SW480^Wild^ cells for heterochromatin associated marks H3K9me3 and H3K27me3, which were significantly altered as determined by immunoblotting. Our data demonstrates a significant loss of these epigenetic marks in the SW480*^PCAT2^*^-KI^ cells sequenced at week 110. We focused our analysis on chromosomes 8 and 4, which harbors our gene of interest *PCAT2* that localize ectopic CENP-A. While we found, in SW480*^PCAT2^*^-KI^ cells, H3K27me3 signatures are significantly lost genome-wide compared to the H3K9me3, chromosome 4 that harbors Trans*PCAT2* gene lost both these signatures (Figure 3A-B and Figure S3A-B). As a representation, we showed a significant loss of H3K27me3 signature in the chromosome 8 in between the known CENP-A ectopic locus 8q24 and 8q21 cytoband with H3K27me3 has more pronounced loss (Figure 3B). In 4q31 locus of SW480*^PCAT2^*^-KI^ cells 93% and 69% of H3K27me3 and H3K9me3, respectively, were lost compared to the SW480^Wild^ cells with only 6% and 25% of peaks were retained. About 2% and 6% of H3K27me3 and H3K9me3 peaks were gained at the 4q31 locus of SW480*^PCAT2^*^-KI^ cells, respectively. Peak distribution analysis showed no significant proportion of peaks were redistributed among the genomic elements, with a light shift in H3K9me3 peaks towards intronic regions (Figure 3C).

**Figure 3.**
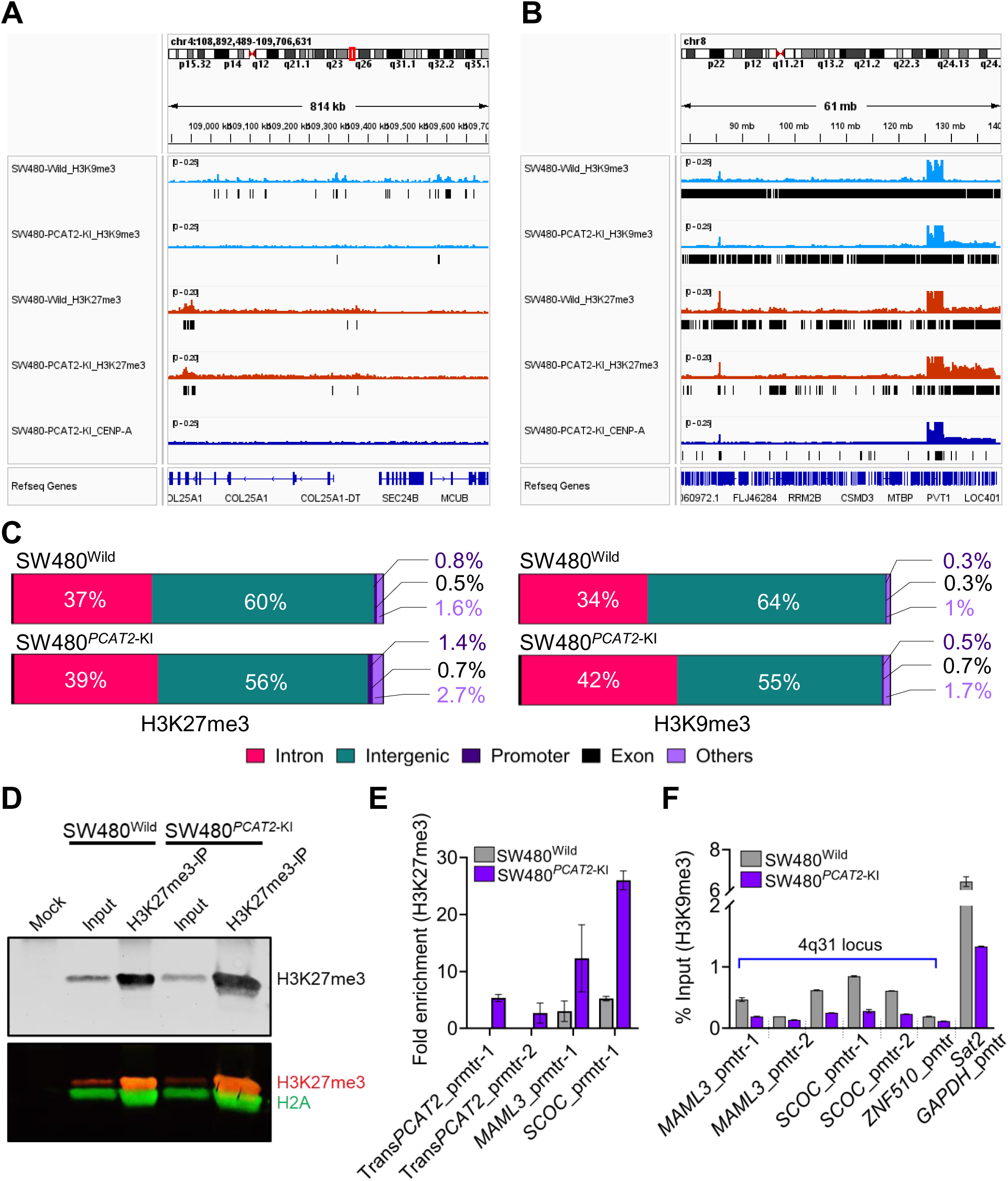
Repressive epigenetic marks silence Trans*PCAT2* gene expression. (A) Representative IGV browser screenshot of chromosome 4 showing significant loss of H3K9me3 consensus peaks are lost in SW480*^PCAT2^*^-KI^ cells compared to SW480^Wild^ cells (blue tracks), while some of the H3K27me3 peaks are retained (red tracks). **(B)** Representative IGV browser screenshot of chromosome 8q locus showing H3K9me3 and H3K27me3 signals are strongly retained at the known CENP-A localizing spots 8q21 and 8q24 between SW480^Wild^ and SW480*^PCAT2^*^-KI^ cells, in contrast to their loss at other sites. **(C)** Peak distribution analysis showed no significant proportion of peaks were redistributed among the genomic elements in SW480*^PCAT2^*^-KI^ cells compared to SW480^Wild^ cells. **(D)** A representative image of immunoprecipitation of chromatin-bound histones probed for H3K27me3 vs. mock IP from SW480^Wild^ and SW480*^PCAT2^*^-KI^ cells. **(E)** Histogram of ChIP-qPCR results shows the level of H3K27me3 signal enrichment at Trans*PCAT2*, *MAML3*, and *SCOC* gene promoters in SW480*^PCAT2^*^-KI^ cells compared to SW480^Wild^ cells. **(F)** Histogram of ChIP-qPCR results showing the levels of H3K9me3 signal enrichment at *MAML3*, *SCOC*, *ZNF510*, *GAPDH* gene promoters, and pericentromeric Sat2 DNA in SW480*^PCAT2^*^-KI^ cells compared to SW480^Wild^ cells.

Furthermore, to determine if the epigenetic changes are targeted to the promoter of the Trans*PCAT2* and neighboring genes at the 4q31 locus of SW480*^PCAT2^*^-KI^ cells we performed ChIP-qPCR analysis. We tested for enrichment of H3K27me3 and H3K9me3 in the Trans*PCAT2* gene carrying SW480*^PCAT2^*^-KI^ cells compared to SW480^Wild^ cells (Figure 3D). Notably, the Trans*PCAT2* cassette is located upstream of *MAML3* and downstream of *SCOC* (Figure 2A). As predicted, we found a significant enrichment of H3K27me3 on the promoter region near the start site of Trans*PCAT2, MAML3,* and *SCOC* genes in the SW480*^PCAT2^*^-KI^ cells (Figure 3E), which also corroborates our IP-western results (Figure 2C-D). However, in contrast to the H3K27me3 mark, we observed a significant depletion of the H3K9me3 mark at these three gene promoters of the SW480*^PCAT2^*^-KI^ cells (Figure 3F). As a control sites we assessed H3K9me3 enrichment on *ZNF510* and *GAPDH* promoters, as well as pericentromeric satellite II (Sat2) DNA repeats. Surprisingly, we found a significant reduction of the H3K9me3 mark at these three sites in SW480*^PCAT2^*^-KI^ cells compared to SW480^Wild^ cells, with a five-fold decrease in pericentromeric Sat2 DNA (Figure 3F). H3K9me3 mark present specifically at Sat2 DNA of chromosome 4 also showed a significant decrease in our ChIP-seq analysis (Figure S3A). The observations suggest a significant impact on the H3K9me3 and H3K27me3 mark globally in the SW480*^PCAT2^*^-KI^ cells with H3K27me3 being more pronounced change. These results collectively suggest that the introduction of the *PCAT2* lncRNA gene to a non-native site and its subsequent overexpression dramatically altered epigenetic signatures and gene expression patterns in SW480*^PCAT2^*^-KI^ cells.

### Trans*PCAT2* gene activity is suppressed by DNA hypermethylation in addition to repressive histone marks

DNA methylation (5mC) is a well-established epigenetic modification crucial in regulating gene expression and genome stability. Typically, DNA methylation at promoter regions is associated with gene repression. In contrast, methylation in the genic regions can either promote or inhibit gene expression, depending on its location and context^48^. Therefore, we were curious to investigate the DNA methylation patterns in Trans*PCAT2* knock-in SW480*^PCAT2^*^-KI^ cells compared to SW480^Wild^ cells. Interestingly, our analysis revealed a genome-wide DNA hypermethylation signature in the SW480*^PCAT2^*^-KI^ compared to SW480^Wild^ cells (Figure S4). While 50% promoters and exonic methylation sites are lost, intergenic and intronic regions exhibit significant DNA-methylation signatures in the SW480*^PCAT2^*^-KI^ cells (Figure 4A). Interestingly, the 4q31 locus of SW480*^PCAT2^*^-KI^ cells carrying Trans*PCAT2* gene knock-in acquired four-fold higher DNA methylation signature than the SW480^Wild^ cells at week 90 (Figure 4B). Notably, the Trans*PCAT2* gene cassette region is widely hypermethylated in the SW480*^PCAT2^*^-KI^ cells (Figure 4C), indicating that the Trans*PCAT2* gene is silenced by both DNA methylation and histone repressive marks (Figure 3 & 4). Overall, the findings reveal that, although DNA methylation has increased globally, the distribution pattern of methylation is uneven. In other words, not all existing methylation sites are hypermethylated; instead, new methylation sites have emerged, while others have been lost. These findings may also be attributed to the dysregulated expression of DNA methyltransferase *DMNTs* and demethylases *TET*, *APOBEC*, and *XRCC* family genes in the Trans*PCAT2* knock-in SW480*^PCAT2^*^-KI^ cells^32^. Our findings suggest that the oncogenic lncRNA *PCAT2* transgene on a naïve chromosome is epigenetically silenced by both DNA methylation and histone repressive marks. Also, introducing a non-native gene to a naïve chromosome site and its overexpression drastically altered the methylation profile of the entire genome.

**Figure 4.**
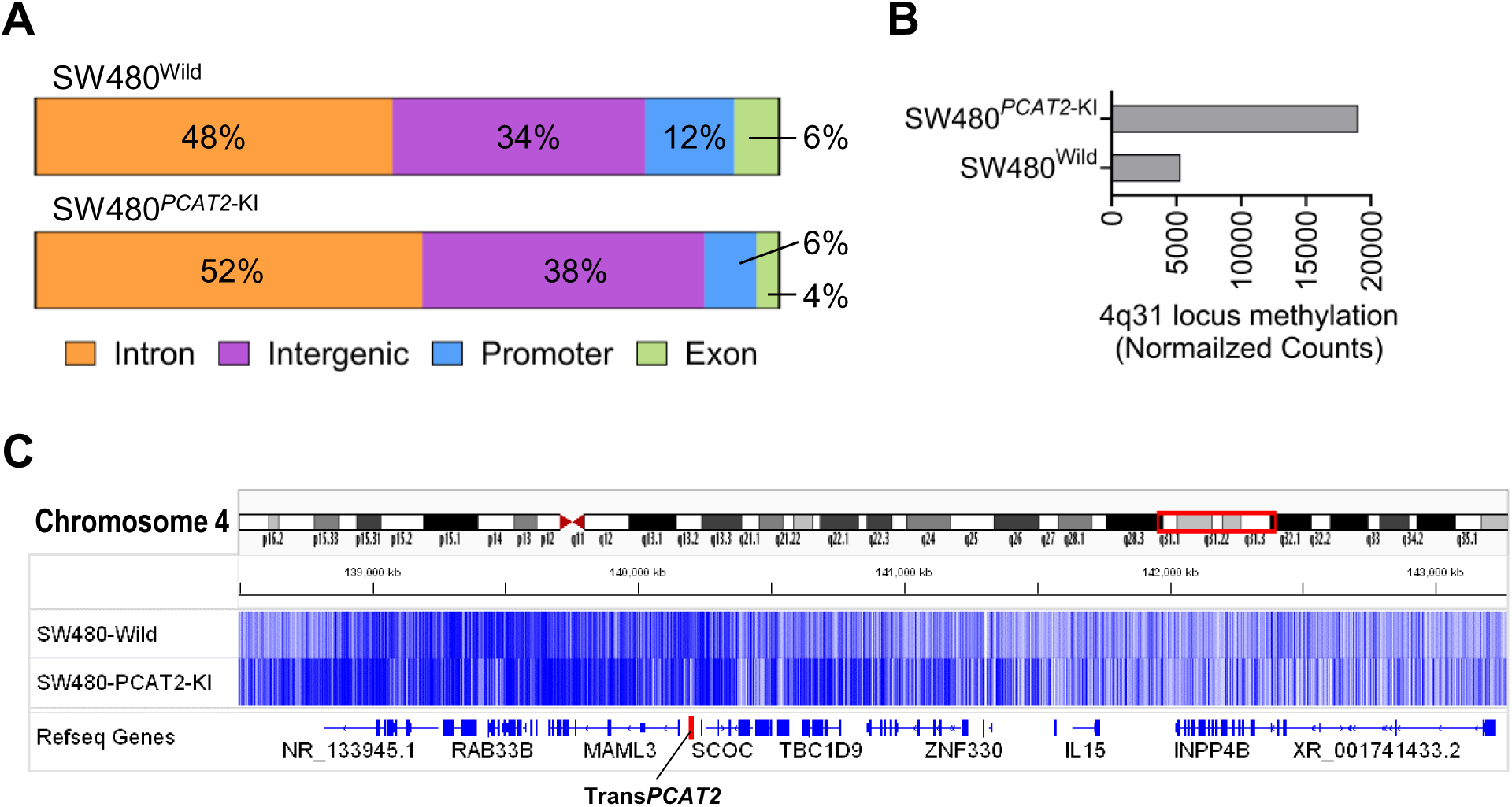
Global methylation profile of Trans*PCAT2* knock-in cells is drastically altered. **(A)** Pie chart shows the percentage of DNA methylation signatures on different genomic regions between SW480^Wild^ and SW480*^PCAT2^*^-KI^ cells. Promoter and exonic regions were hypomethylated, while intergenic and intronic regions were hypermethylated in SW480*^PCAT2^*^-KI^ cells. **(B)** Bar graph showing the normalized counts of methylation signals from 4q31 locus of SW480*^PCAT2^*^-KI^ cells is four-fold higher than SW480^Wild^ cells. **(C)** The genome browser screenshot of the 4q31 locus shows that the Trans*PCAT2* knock-in site at the 4q31.1 locus of SW480*^PCAT2^*^-KI^ cells has a significant hypermethylation signature compared to SW480^Wild^ cells.

### Multiple epigenetic silencing mechanisms prevent the reactivation of the Trans*PCAT2* gene

Next, we were interested in reactivating the expression of the Trans*PCAT2* gene by using heterochromatin-altering drugs in the SW480*^PCAT2^*^-KI^ cells. Forskolin promotes the PKA-CREB- dependent signaling pathway through CREB phosphorylation by PKA. Activated CREB binding to CRE sites enhances the recruitment of other transcriptional machinery, leading to activation of gene promoters and increased expression^49^. Forskolin is also shown to reactivate silenced CMV promoters^50^, and the engineered Trans*PCAT2* genes are driven by CMV promoters. Therefore, we hypothesized that reactivating Trans*PCAT2* expression could repopulate the ectopic CENP-A at the 4q31 locus. To achieve this, we treated the SW480*^PCAT2^*^-KI^ cells with Forskolin to reactivate the CMV promoter driving the expression of the Trans*PCAT2* genes. The Forskolin treatment did not reactivate Trans*PCAT2* and increase the ectopic CENP-A level at the 4q31 locus (Figure 5A). The CENP-A levels at the 8q24 locus remained unchanged upon Forskolin treatment. This result suggests that the Trans*PCAT2* gene may be strongly silenced through multiple epigenetic mechanisms that prevents its transcriptional activation^51^.

**Figure 5.**
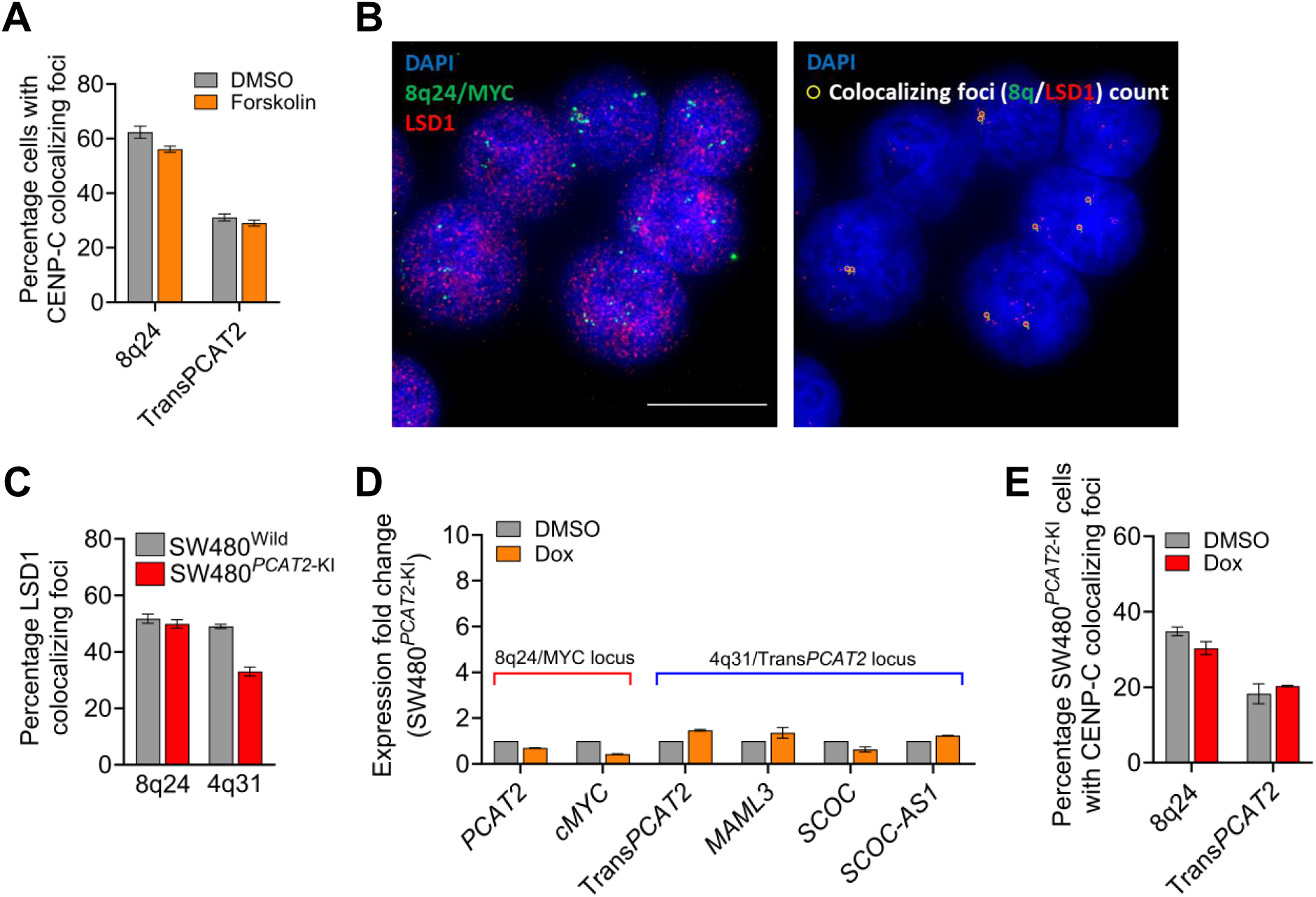
Epigenetic silencing mechanisms prevent reactivation of TransPCAT2. **(A)** Histogram shows the levels of CENP-C colocalized at 8q24 and 4q31 locus after Forskolin treatment compared to DMSO controls in SW480*^PCAT2^*^-KI^ cells. **(B)** Representative immunofluorescence image shows the LSD1 colocalization at 8q24 locus in SW480^Wild^ cells. The scale bar represents 10 µm. **(C)** Histogram shows the levels of LSD1 colocalized at 8q24 and 4q31 locus in SW480*^PCAT2^*^-KI^ cells compared to SW480^Wild^ cells. **(D)** Histogram shows the expression levels of *PCAT2* and *cMYC* genes at the 8q24 locus and Trans*PCAT2*, *MAML3*, *SCOC*, and *SCOC-AS1* genes at the 4q31 locus of SW480*^PCAT2^*^-KI^ cells compared to SW480^Wild^ cells after doxorubicin treatment. **(D)** Histogram shows the percentage of SW480*^PCAT2^*^-KI^ cells that colocalize CENP-C at 8q24 and Trans*PCAT2* foci with doxorubicin treatment compared to DMSO control. No significant change was observed in both the foci after doxorubicin treatment.

### Drug-induced heterochromatin disruption does not activate Trans*PCAT2* gene transcription

Finally, we were curious to investigate the occupancy of chromatin modifiers that scaffold for assembling transcription factors or silencers to activate or repress transcription. For this, we tested for occupancy of LSD1 (Lysine-specific demethylase 1), also known as KDM1A, which targets mono- or di-methylated histone H3K4 and H3K9^52^. LSD1 inhibits the formation of heterochromatin by demethylating H3K9^53^. The removal of H3K9 methylation allows heterochromatin to open, which facilitates gene expression. Surprisingly, we observed a reduction in LSD1 localization at the Trans*PCAT2* gene region on the 4q31 locus in SW480*^PCAT2^*^-KI^ cells at week 90 compared to SW480^Wild^ cells (Figure 5B & C). The diminished localization of LSD1 at the Trans*PCAT2* gene region further supports our previous observation of heterochromatin formation (Figure 3). Since the 4q31 locus displays heterochromatin formation, we next sought to employ a drug-mediated heterochromatin disruption and DNA demethylation to reactivate the Trans*PCAT2* gene. Doxorubicin inhibits the catalytic activity of DNMT1 through DNA intercalation and alter chromatin structure by affecting nucleosome turnover, influencing the binding of architectural proteins like HMGB1 and linker histone H1, and potentially disrupting spatial organization of the genome^54–57^. Therefore, we treated the SW480*^PCAT2^*^-KI^ cells with doxorubicin for 48 hours and monitored the transcript levels of 4q31 and 8q24-derived genes to indirectly access the disruption of heterochromatin and activation of Trans*PCAT2* gene. We observed that doxorubicin treatment slightly increased the expression of the Trans*PCAT2* gene and a few neighboring genes at the 4q31 locus (Figure 5D). Since dox treatment is not targeted to 4q31 locus, we also tested the gene expression levels at 8q24 locus. The expression of the innate copy of *PCAT2* and *cMYC* at the 8q24 locus was slightly decreased (Figure 5D). Overall, dox-treatment to SW480*^PCAT2^*^-KI^ cells did not significantly alter the ectopic CENP-A levels (Figure 5E). Thus, we concluded that the drug-induced activation strategies did not reactivate the Trans*PCAT2* locus in a shorter treatment time to alter the ectopic CENP-A occupancy significantly. This could be due to the Trans*PCAT2* cassette being inserted into a lncRNA deprived low-transcribing locus, unlike the lncRNA rich 8q24 locus which helps keep the chromatin open and highly active for a shorter drug treatment period, thereby reactivating the locus. Overall, our results suggest that multiple layers of epigenetic controls, as a defensive mechanism, may suppress the Trans*PCAT2* gene at the 4q31 locus in SW480*^PCAT2^*^-KI^ cells.

## Discussion

Many types of cancer display overexpression of CENP-A and subsequent ectopic localization. Most CENP-A ectopic sites are high-transcription sites and subtelomeric and telomeric regions within the genome^28^. However, the mechanism responsible for CENP-A ectopic localization remains unclear. Our previous report demonstrated that an oncogenic lncRNA *PCAT2* transcription can recruit CENP-A to a naïve chromosome locus, leading to a significant ectopic site^32^. Additionally, *PCAT2* was found to interact physically with CENP-A, allowing it to mimic centromeric transcription and recruit excess CENP-A to its transcribing locus through a non-canonical chaperone pathway. However, it remains unclear how stable this induced ectopic site is in establishing epigenetic memory and whether it can persist in creating a stable neocentromere. Answering these questions could provide valuable insights into the development and evolution of neocentromeres through lncRNA-mediated epigenetic dysregulations. Therefore, we tracked the artificially persuaded CENP-A ectopic site by the lncRNA Trans*PCAT2* gene at the 4q31 locus of a colon cancer cell line for over two years.

Following an initial spike in ectopic CENP-A level at the Trans*PCAT2* gene region^32^, for about 20 weeks, the CENP-A level steadily declined and eventually returned to its basal level by week 40 and remained stable. Trans*PCAT2* expression was much higher in the early weeks before dropping dramatically, which coincided with ectopic CENP-A level at the 4q31 locus. Notably, nearby genes were silenced while Trans*PCAT2* levels were significantly elevated. However, the 4q31 gene expression levels recovered to normal while the Trans*PCAT2* transcript level reduced over time. This shows that introducing Trans*PCAT2* to a non-native location on a naïve chromosome locus affects the normal function of that genomic region. The change in gene expression at the 4q31 locus is likely due to altered *cis*-regulation, such as promoter-enhancer interactions and dysregulation of key epigenetic regulators due to Trans*PCAT2* overexpression. Furthermore, enrichment of H3K27me3, DNA hypermethylation, and reduced LSD1 localization at the Trans*PCAT2* gene region in SW480*^PCAT2^*^-KI^ cells indicate an extensive epigenetic rewiring occurring at the 4q31 locus. H3K27me3 is often associated with the repressive function of lncRNA genes, and the reaccumulation of this mark is frequently observed in genes with DNA hypermethylation in cancer cells^58–60^. Therefore, the enrichment of H3K27me3 and DNA methylation at the Trans*PCAT2* gene region could indicate a robust epigenetic silencing mechanism. However, it remains unclear whether the epigenetic silencing of the Trans*PCAT2* gene is a result of loss of *cis*-regulatory elements that were initially present at this site before introducing the transgene cassette or if it is a failsafe mechanism implemented by cells to control *PCAT2* transcript level.

CENP-A deposition at the centromeres is a transcription-mediated process, and ectopic CENP-A is seen at high-transcription turnover sites. Therefore, reactivating the Trans*PCAT2* promoter for transcription by demethylating or redistributing euchromatin/heterochromatin boundaries could restore the ectopic CENP-A to the 4q31 locus. However, drug treatments were ineffective in significantly transcriptionally reactivating Trans*PCAT2,* and no new recruitment of CENP-A was observed. This lack of reactivation can be attributed to multiple layers of epigenetic silencing mechanisms, including H3K36me2, H3K79me1, H4K20me1, etc., in addition to the tested H3K9me3 and H3K27me3 modifications^61^. Additionally, the active or inadequate recruitment of epigenetic regulatory factors such as EZH2, SET family, MLL family, GLP, CLL8, NSD1, etc., may contribute to the continued inactivation of the Trans*PCAT2* gene promoter^61^. Furthermore, the absence of a mammalian enhancer element, repositioning of the Trans*PCAT2* gene to the nuclear periphery, and the loss of distal activating *cis*-elements concerning to this transgene could also significantly impact its transcriptional capacity over time^62–66^. Although CpG methylation has been shown to silence specific genes in cancer cells, evidence for targeted gene silencing *via* histone modification is scarce^67^. One theory is that because the SW480 colon cancer cells employed in this work already had five copies of the 8q24 region encompassing the *PCAT2* gene, the cells may not require more *PCAT2* transcript. However, it is unclear whether all copies of PCAT2 genes are actively transcribed in these cells. Moreover, *PCAT2* transcription recruits CENP-A to its location and establishes an ectopic domain for CENP-A, which is eventually expected to develop into a neo-centromere over time. Such neocentromere formation, however, may be harmful to the individual chromosome and may not be a favorable selection for cancer cells. This is also evident from our ChIP sequencing results, which show that the CENP-A peaks that spiked in the initial period of the Trans*PCAT2* knock-in vanished at a later time. It is only specific aneuploidies that are common, and they are frequently cancer-specific^68^. Given that 4q deletions are uncommon^69^, but have been observed in microsatellite-stable colon tumors^70^, it is likely that SW480 cells do not favor 4q31 fragility *via* the ectopic CENP-A deposition pathway.

However, additional research is required to understand why certain aneuploidies are prevalent and specific to cancer and investigate the mechanisms underlying the stability of neocentromeres. Nonetheless, the combined effect of multiple lncRNAs may create a more permissive environment (Figure 6), characterized by a broad segment of open chromatin, high transcription rate, frequent and sustained R-loop formation, enrichment of non-B-DNA structures like G- quadruplex DNA, and the presence of prerequisite factors for CENP-A-non-canonical histone chaperones at the *PCAT2* native 8q24 locus^28,29,71,72^. Our previous observation that ectopic CENP-A levels are significantly lower at the innate 8q24 locus compared to the translocated 8q24 locus^32^, suggests that *cis*-regulation plays a significant role in determining the transcriptional activity of the locus for CENP-A recruitment. Additionally, the combinatorial inhibition of various 8q24 locus lncRNAs, along with *PCAT2*, significantly suppressed the localization of CENP-A^32^, highlighting the collective function of genes from a chromosomal locus in the retention of ectopic CENP-A. This motivates further exploration of the hypothesis that an entire 8q24 locus can be translocated to a naïve chromosome in 8q24 duplication null cells, and studying the effective recruitment of CENP-A to this locus. Alternatively, CRISPR-KRAB-mediated suppression of all innate 8q24 locus lncRNAs, as demonstrated for *PCAT2* lncRNA^32^, could be employed to investigate whether and for how long a large ectopic CENP-A domain can be sustained without active transcription. Further investigation into whether the combined action of 8q24 locus lncRNAs or a single, yet competent, lncRNA transcription is sufficient for the active recruitment and long- term maintenance of ectopic CENP-A may be possible by simply replacing the CMV promoter with a mammalian promoter or the native *PCAT2* gene promoter. Therefore, this ongoing transitional work marks a significant step towards unraveling the mechanisms of lncRNA- mediated epigenetic regulation and chromosome instability in cancer cells. Furthermore, this study emphasizes the importance of considering global epigenetic signatures when investigating loss- or stable transgene-mediated gain-of-function of lncRNAs.

**Figure 6.**
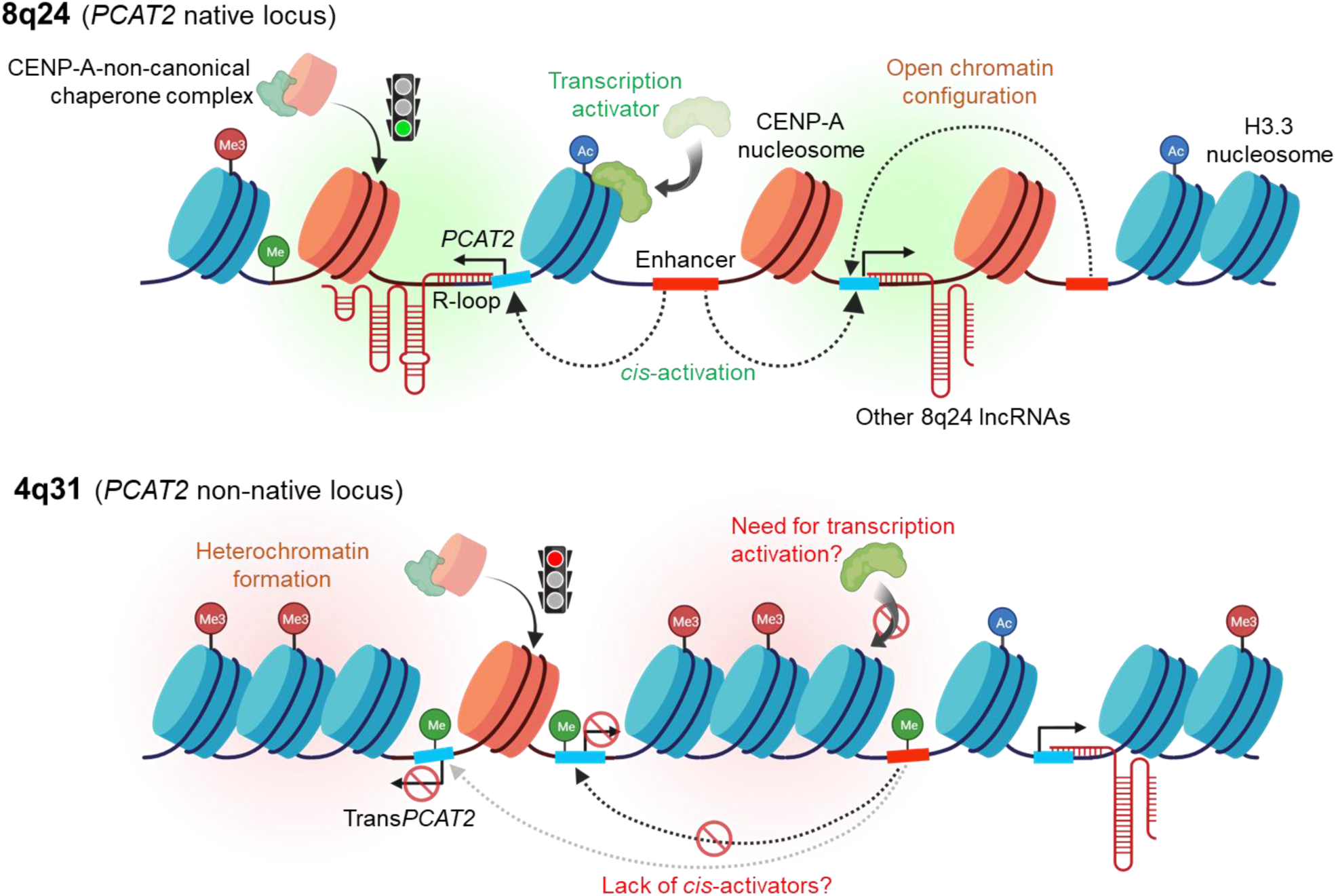
Epigenetic silencing of an oncogenic lncRNA transgene cassette. A graphical representation of our model illustrates the cellular fail-safe mechanism through various epigenetic modifications. The active transcription of an oncogenic lncRNA *PCAT2*, expressed from its native locus on the chromosome 8q24 region, can recruit and maintain CENP-A at its transcribing locus. This recruitment may be facilitated by multiple mechanisms, including persistent transcriptional activation by *cis*- and *trans*-activating factors, an open chromatin environment created by neighboring lncRNA transcription, formation of R-loops and non-B-DNA structures, and factors that enable the binding of a CENP-A-non-canonical chaperone complex for CENP-A deposition. However, when the same *PCAT2* lncRNA gene is introduced to a non-native chromosome locus, the absence of *cis*- and *trans*-activating factors and the lack of an open chromatin confirmation weaken the sustained expression of the *PCAT2* gene. This loss of active and sustained transcription of *PCAT2* at the naïve locus ultimately leads to the failure to recruit new CENP-A to its transgene locus. Additionally, cellular fail-safe epigenetic silencing mechanisms, such as DNA methylation and histone modifications on the H3.3 nucleosomes, come into play to prevent the reactivation of *PCAT2* transcription and CENP-A recruitment. Therefore, we propose that even cancerous cells may have evolved several epigenetic pathways to silence deleterious events and prevent excessive chromosome damage, ensuring their survival.

We report that the induced epigenetic memory generated by oncogenic lncRNA, which results in the misplacement of CENP-A to an ectopic small transgene cassette, is transient and does not lead to sustained transcriptional activity when compared to the much larger innate locus that contains a complete gene regulatory element. Importantly, the maintenance of CENP-A at centromeres requires active transcription of centromere DNA and may depend on various factors, including CENP-C, CENP-B, and active marks such as H3K4me2 domains. Consequently, a lack of these factors could render this transgene cassette a weak and transient ectopic CENP-A domain. In response to unfamiliar and detrimental chromosome-damaging events, we hypothesize that cells may activate a protective mechanism involving epigenetic silencing around a target locus. This mechanism helps prevent potential detrimental consequences, which may vary depending on the specific cell type. Consequently, we are committed to further investigating the role of lncRNAs in epigenetic rewiring, the preferential enrichment of specific aneuploidies, and the underlying epigenetic regulatory mechanisms that drive these events in the positive selection of cancer cells.

## Methods

### Cell culture and Treatment conditions

The SW480 (SW480^Wild^ and SW480*^PCAT2^*^-KI^) human colon cancer cells were cultured in RPMI 1640 medium (catalog no. 11875093, Gibco, USA) with glutathione, 10% fetal bovine serum (catalog no. S11050H, Atlanta Biologicals, USA), and 1x penicillin-streptomycin solution (catalog no. 15140122, Gibco, USA) at a temperature of 37°C and 5% CO_2_ in a humidified incubator. In the case of CMV promoter reactivation experiments, the ∼5 million cells were seeded 24 hours before drug treatment. Next, 1μM Forskolin (catalog no. F3917, Sigma-Aldrich, USA) was added directly to the culture media and incubated for three days before harvesting the cells for downstream analysis. For heterochromatin alteration by Doxorubicin, similar to Forskolin treatment, cells were seeded 24 hours before inhibition treatments. Doxorubicin (catalog no. D1515, Sigma-Aldrich, USA) (250 nM) was added directly to the culture media. The cells were harvested after 48 hours for downstream experiments.

### RNA isolation

RNAs were extracted using TRIzol reagent (catalog no. 15596026, Ambion, USA) according to the manufacturer’s protocol. For 1 ml of TRIzol-containing sample, 200 μl of chloroform was added, gently vortexed, and incubated for 4 min at room temperature. After centrifugation at 12,000 rpm for 15 min at 4°C, the clear upper phase was transferred to a fresh tube. To the clear phase, 500 μl of isopropanol (catalog no. 534021, Sigma-Aldrich) was added and incubated at −20°C for 30 min. The samples were centrifuged again, and the RNA pellet was washed with 1 ml of cold 75% ethanol and resuspended in diethyl pyrocarbonate ultrapure water (catalog no. RGF-3050, KD Medical, USA). Isolated RNA was quantified using a DeNovix DS-11 spectrophotometer (DeNovix Inc., USA)

### Complementary DNA synthesis and Real-time qRT PCR

Complementary DNAs (cDNA) were synthesized using 2 μg with a SuperScript III Reverse Transcriptase SuperScript™ III First-Strand Synthesis SuperMix kit (catalog no. 11752050, Invitrogen, USA) following the manufacturer’s protocol. The cDNAs were diluted in nuclease-free water and stored at −20 °C until use.

cDNA samples were prepared using GreenLink™ Blue qPCR Mastermix (catalog no. 16- 2200, BioLink laboratories, USA) following the manufacturer’s protocol. The quantitative reverse transcription PCR (qRT-PCR) was run on the QuantStudio 3 Real-time PCR System (Applied Biosystems, USA), and relative quantification was performed using the 2^−ΔΔCT^ method. All reactions were performed in 20 μl volume, and triplicates with no template controls were performed to rule out nonspecific amplification. *GAPDH* served as an internal control. Melting curve analysis was performed for all the primer sets to check the specificity of the primers. Primer sequences used in this study are listed in supplement table S1.

### RNA sequencing

Total RNA was isolated from SW480*^PCAT2^*^-KI^ colon cancer cells. RNA quality was assessed using the 2100 Bioanalyzer (Agilent, USA). Sequencing libraries were prepared from the total RNA samples and sequenced G4™ Sequencing Platform (Singular Genomics, USA) in paired-end mode. Raw reads underwent preprocessing to eliminate low-quality bases and adapter sequences with Trimmomatic. The trimmed reads were then mapped to the human genome (GRCh38) using Tophat2 v2.0.8. To determine differential expression, read counts for genes were generated using Subread 2.0.6 against Ensembl release 103 annotations. Differential expression analysis was performed with DESeq2, reporting results at a significance threshold of a false discovery rate-adjusted P value of <0.01. Transcript abundance was quantified in RPKM (reads per kilobase of transcript per million reads mapped).

### Whole-genome sequencing and Bisulfite sequencing

High molecular weight DNA from SW480 (SW480^Wild^ and SW480*^PCAT2^*^-KI^) colon cancer cells were isolated using PacBio® Nanobind® HMW DNA extraction kits (catalog no. 102-762- 700, PacBio, USA) following the manufacturer’s protocol. The quality of the DNA was tested using 2100 Bioanalyzer (Agilent, USA) and subjected to long-read sequencing *via* Single-Molecule Real-Time (SMRT) technology on a PacBio Sequel IIe Sequencer (PacBio, USA) for whole-genome sequencing. The sample was mapped, and variants were called using the DRAGEN v4.0.3 pipeline (Illumina, USA). Using a blast search algorithm against a custom reference with the insert plasmid sequence, the insert sequence was identified from the whole-genome sequencing reads and mapped.

For methylation sequencing, following library preparation, the pooled bisulfite-modified samples underwent sequencing on a NovaSeq 6000 S1 (Illumina, USA), run using paired-end sequencing mode. Subsequently, the sequences were mapped, and methylation sites were identified utilizing the DRAGEN v4.0.3 pipeline software (Illumina, USA) against the reference genome hg38. The Ensembl genome features were employed to annotate the methylated sites. Differential analysis for the identified methylation sites was conducted using the MethylKit function to calculate DiffMeth^73^. Differentially methylated sites were determined by a minimum 15% change in methylation with a *q*-value of < 0.01 and a coverage of at least 10X.

Methylation calls were extracted from the bam files using the bismark_methylation_extractor from Bismark, a bisulfite-aware aligner with the --comprehensive and --bedGraph options to obtain strand-specific methylation counts in CpG context. Methylation data were filtered to include CpG sites within the 4q31 locus (chr4:145,000,000–146,000,000). For each sample, methylated and unmethylated read counts were extracted from Bismark “.cov” files and summed across the locus to obtain total counts per condition. These counts were used to assess global methylation status across the region. Stacked bar plots comparing methylated and unmethylated read counts in the 4q31 locus were generated using R (ggplot2).

### Chromatin immunoprecipitation and Immunoblotting

Five flasks of cells were grown in the T175 flasks (catalog no. 83.3912, Sarstedt, USA) and harvested using trypsin (catalog no. 25200056, Gibco, USA). The cells were washed with PBS and cold PBST (0.1% Tween). After spinning down, the cell pellet was washed with cold TM2 buffer containing NP-40 (catalog no. FNN0021, Invitrogen, USA) and resuspended in 0.1 M TE. The cell pellet was then treated with MNase (4 U/ml) (catalog no. 9013-53-0, Millipore Sigma, USA) for 15 min in a 37°C water bath, and chromatin was extracted overnight using a low-salt buffer with 1X cOmplete Protease Inhibitor Cocktail (catalog no. 11697498001, Millipore Sigma, USA) at 4°C. Inputs were collected at this point and stored at −20°C. The supernatant collected was incubated with our custom CENP-A antibody (rabbit polyclonal, Epitope: C- TPGPSRRGPSLGA) overnight at 4°C in a rotator. The CENP-A antibody-bound chromatin was pulled down in the low-salt buffer using Dynabeads® Protein G-tagged magnetic and sepharose beads (catalog no. 10003D, Invitrogen, USA).

The precipitated samples were treated with Proteinase K (catalog no. AM2548, Ambion, USA) at 55°C overnight, and phenol: chloroform: isoamyl alcohol (catalog no. P3803, Sigma-Aldrich, USA) was used to isolate DNA for ChIP-qPCR^32^. In the case of immunoblotting, the ChIP samples were mixed with 4X Laemmli sample buffer (catalog no. 1610747, Bio-Rad, USA), denatured for 10 min at 95°C, and incubated on ice for 2 min. The samples were resolved in 4 to 20% Mini-PROTEAN^®^ TGX^TM^ precast gels (catalog no. 4561093, Bio-Rad, USA), with 1X tris-glycine SDS running buffer (catalog no. RGC-3390, KD Medical, USA), and transferred to nitrocellulose membrane using Trans-Blot^®^ Turbo^TM^ Mini Transfer Packs (catalog no. 1704158, Bio-Rad, USA). The membrane was blocked in 1:1 Odyssey blocking buffer (catalog no. 92740000, LI-COR, USA) at room temperature for 30 min and incubated with primary antibody diluted in 1:1 blocking buffer and 1X PBS complemented with 0.1% Tween 20 on a rocker for overnight at 4°C. After three washes in 1X PBS, 0.1% Tween 20, the membrane was incubated with the Alexa Fluor (Invitrogen, USA) or IRDye^®^ (LI-COR, USA) conjugated to the secondary antibody diluted in blocking buffer supplemented with 0.1% Tween 20% for 1 hour at room temperature. Finally, the membrane was washed once with 1X PBST and imaged/analyzed using an Odyssey^®^ M Imager and Empiria Studio^®^ Software (LI-COR Biosciences, USA).

All antibodies used in this study are commercially available [CENP-A (catalog no. ab13939; Abcam, USA/catalog no. 07-574-MI; Millipore Sigma, USA), CENP-C (catalog no. PD030, MBL, USA), GAPDH (catalog no. SC20357, Santa Cruz Biotechnology, USA), H3K27ac (catalog no. ab4729; Abcam, USA), H3K27me3 (catalog no. ab192985; Abcam, USA), H3K9me3 (catalog no. ab176916; Abcam, USA), H2A (catalog no. ab18255; Abcam, USA), and H3.3 (catalog no. SC8654, Santa Cruz Biotechnology, USA)].

### ChIP sequencing

Five flasks of SW480*^PCAT2^*^-KI^ colon cancer cells were grown in the T175 flasks and harvested using trypsin. The cells were washed once with PBS and once with cold PBST (0.1% Tween). The pellet was washed with cold TM2 buffer containing NP-40 (catalog no. FNN0021, Invitrogen, USA) and resuspended in 0.1 M TE. The cell pellet was then treated with MNase (4 U/ml) for 15 min in a 37°C water bath, and chromatin was extracted overnight using a low-salt buffer with 1× Protease Inhibitor Cocktail (catalog no. G6521, Promega, USA) at 4°C. The supernatant was incubated overnight at 4°C with appropriate antibodies. The antibody–bound nucleic acids were pulled down using Dynabeads Protein G–tagged magnetic beads (catalog no. 10003D, Invitrogen, USA) in the low-salt buffer. The nucleic acid–bound beads were then subjected to RNase I treatment (NEB, USA) and DNA extraction using the phenol-chloroform protocol. The DNA was sequenced in G4™ Sequencing Platform (Singular Genomics, USA) using paired-end reads. Data were processed, normalized, and analyzed, as previously mentioned ^28,32^.

### Metaphase chromosome preparation

Cells were grown in a T175 flask, and colcemid (catalog no. 10295892001, Millipore Sigma, USA) was added 8 hours before the end of harvest time. The cells were harvested through trypsinization (catalog no. 25200056, Thermo Fisher Scientific, USA). The cells were washed with 10 ml of 1× PBS, then suspended in 6 ml of hypotonic solution and incubated in a water bath at 37°C for 20 minutes. They were fixed using a freshly prepared cold fixative solution (methanol and glacial acetic acid in a 3:1 ratio). After centrifuging at 1500 rpm for 5 minutes at room temperature, the cells were resuspended in 4 ml of fixative and incubated for 15 min at room temperature. The cells were centrifuged again, resuspended in 400 μl of the fixative solution, and stored at 4°C until slide preparation.

### Immunofluorescence and DNA-Fluorescence in situ hybridization

As mentioned in our previous publication^32^, Immunofluorescence and DNA-fluorescence in situ hybridization protocols are followed. Briefly, the metaphase chromosomes were dropped onto the glass slides and air-dried. The slides were immediately incubated in the TEEN buffer [1 mM triethanolamine-HCl (pH 8.5), 0.2 mM Na-EDTA, and 25 mM NaCl], followed by blocking [0.1% Triton X-100 and 0.1% BSA in the TEEN buffer] and antibody incubation steps in KB buffer [10 mM Tris-HCl (pH 7.7), 150 mM NaCl, and 0.1% BSA]. In the case of unfixed cells, the cells were pounded onto the glass slide using a Cytospin^TM^ (Thermo Scientific, USA), and the slides were subjected to the immunofluorescence protocol mentioned above.

For DNA-FISH, the slides were equilibrated in 2X SSC at room temperature and digested with Pepsin (catalog no. P6887, Sigma-Aldrich, USA) in 2X SSC (10 mg/ml; catalog no. 10108057001, Millipore Sigma, USA) at 37°C. The slides were washed in 2X SSC and dehydrated in ethanol series (70, 95, and 100%). Following dehydration, the slides were denatured in 70% formamide in 2X SSC at 80°C and dehydrated again in ethanol series. The DNA-FISH probe was added to the slides, covered with a coverslip, sealed with rubber cement, and incubated in a humidified chamber at 37°C overnight. After washes with 2× SSC containing 50% formamide and 0.2X SSC, the slides were air-dried and mounted using a DAPI-containing mounting solution in the dark. DNA FISH probes tagged with fluorophores were procured commercially from Empire Genomics (USA).

### Imaging and statistical analysis

DNA-FISH and IF slides were imaged at 60X in a DeltaVision^TM^ Elite RT microscopy imaging system (GE Healthcare, USA) with a charge-coupled device camera (CoolSNAP, USA) mounted on an inverted microscope (IX-70, Olympus, USA). The acquired images were deconvolved and analyzed with ImageJ (version 1.51 with Java 1.8.0_172). Proteins colocalizing at DNA-FISH-probed loci were identified using the ImageJ software’s “Colocalization” plugin. All numerical data are presented as the mean ±SD in graphs. The number of loci counted for colocalization is presented with *p*-values in supplement table S2. A two-tailed Student’s t-test was used to calculate the differences between means. Fisher’s exact test was used to analyze the difference between groups using GraphPad Prism software (v10.1.1, GraphPad Software Inc., USA). A *p*-value of <0.05 was considered statistically significant. Illustrative diagrams were created using BioRender (BioRender, USA) and Photoshop CC 2019 (Adobe Inc., USA).

## Acknowledgment

We thank the NCI-CCR core sequencing facility for supporting sequencing whole-genome and bisulfite sequencing and the UNMCC Analytical and Translational Genomics Shared Resource core for supporting RNA and ChIP sequencing. We also acknowledge CSEM members for their advice and feedback.

## Author contributions

G.A. and Y.D. conceptualized the research. G.A. designed and performed experiments, standardized protocols, acquired and analyzed the data, consolidated the results, and wrote the manuscript. S.S. performed ChIP-western experiments, consolidated the results, assisted writing and reviewing the manuscript. Y.D. and G.A. reviewed the data and manuscript. S.B. performed all computational analysis.

## Data availability

The paper and the supplementary materials contain all the data needed to evaluate the conclusions. Sequenced reads have been deposited in the Gene Expression Omnibus (GEO accession GSE249606).

