## Supplemental Figures S1-S4 for "Oncogenic lncRNA transgene transcription modulates epigenetic memory at a naïve chromosomal locus"

### Figure S1

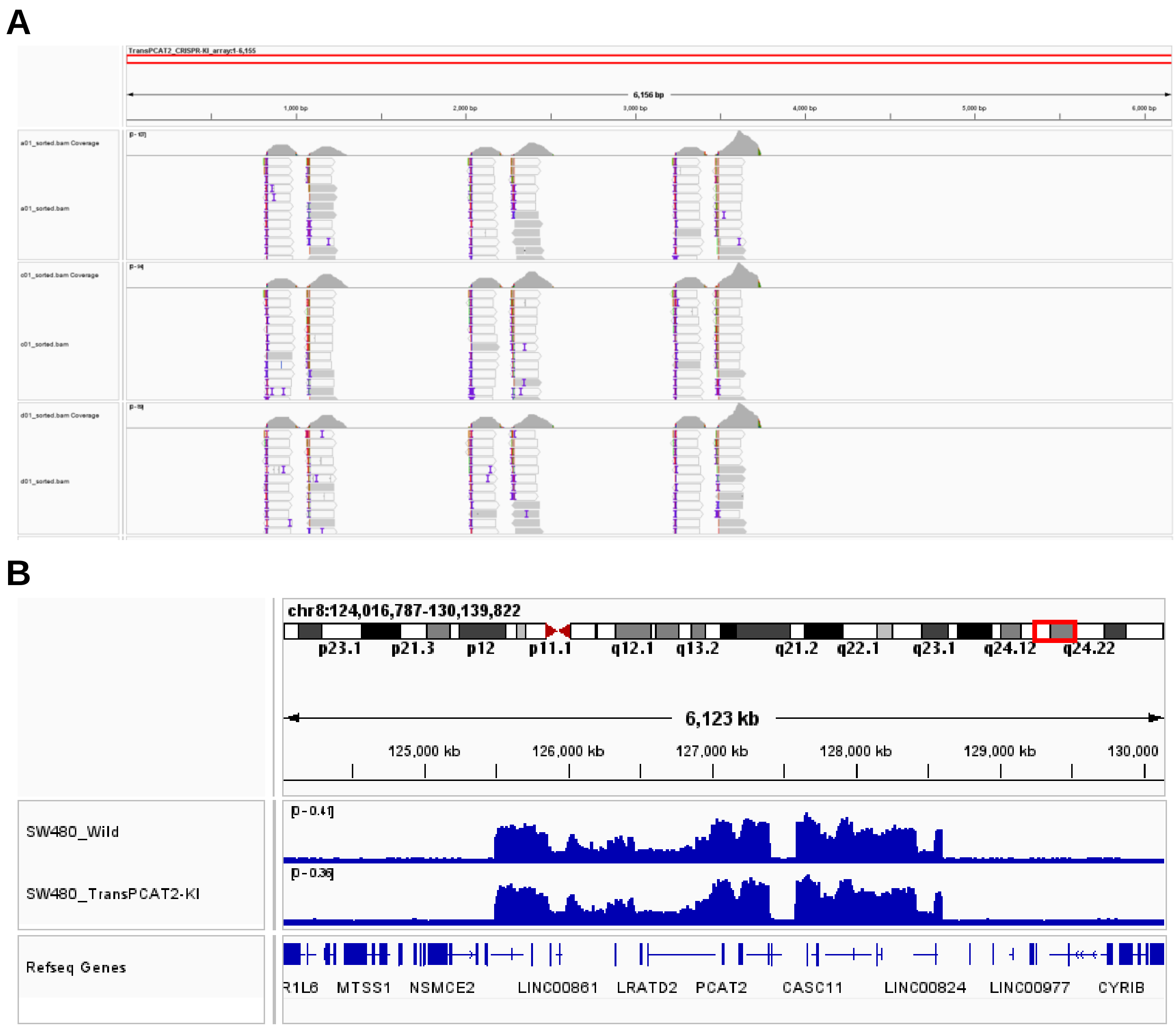


**Figure S1. Confirmation of Trans*PCAT2* gene and CENP-A occupancy in stable cells. A.** IGV browser screenshot of sequence reads for the TransPCAT2 gene at the CRISPR knock-in site in the chromosome 4q31 locus of SW480$^{\mathit{PCAT2}\text{-KI}}$ colon cancer stable cells. The long-read sequences are trimmed into shorter sequences and blasted against a custom-built chromosome 4 sequence assembly using the Trans*PCAT2* array sequence. **B.** IGV browser screenshot of ChIP-sequencing using anti-CENP-A, showing no change in CENP-A ectopic occupancy at the 8q24 locus between SW480 wild-type and SW480$^{\mathit{PCAT2}\text{-KI}}$ cells.

### Figure S2

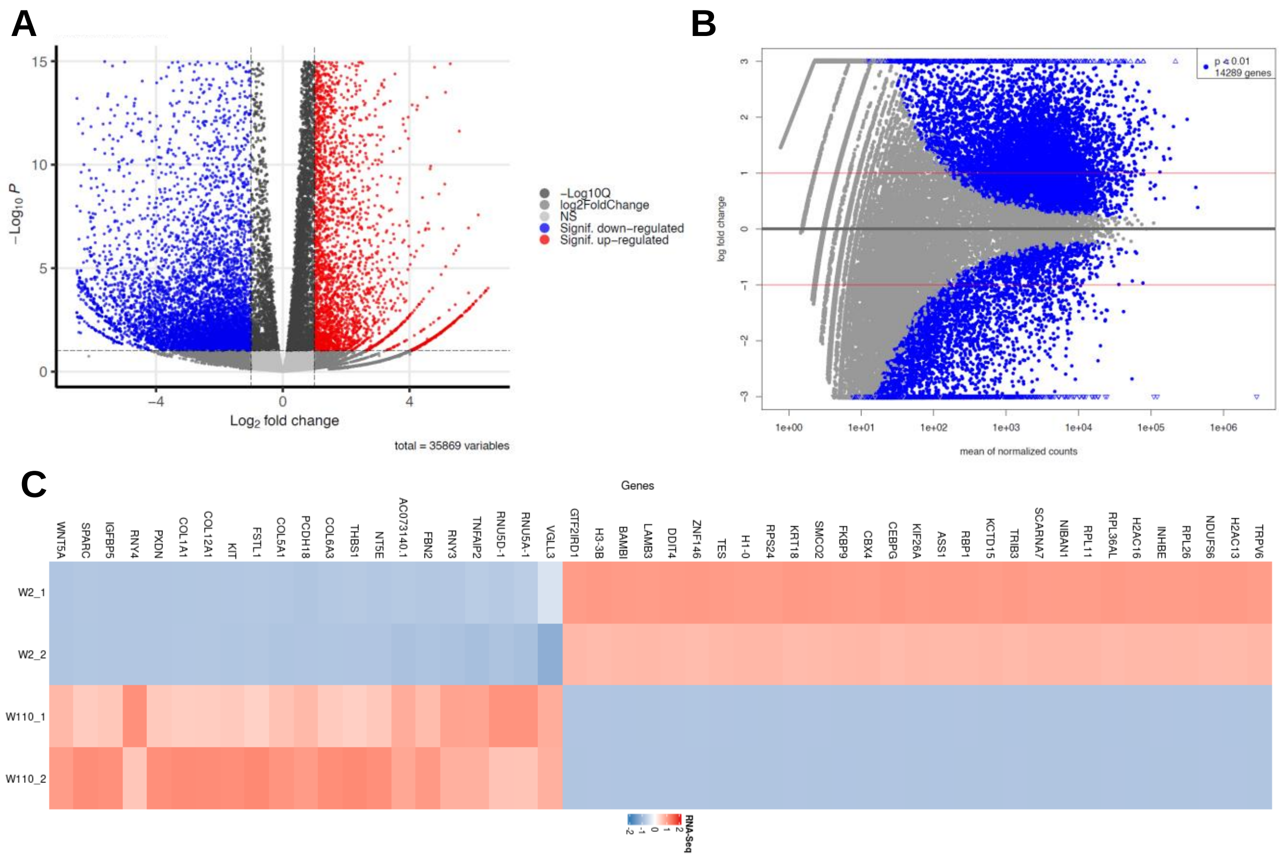


**Figure S2. Differential gene expression signature. A.** Volcano plot showing the expression pattern of genes in SW480$^{PCAT2\text{-KI}}$ cells sequenced at week 2 (W2) and week 110 (W110). Over 35000 genes are differentially expressed between two time periods. **B.** An MA plot visualizes the relationships between the log ratio and mean values of genes in these two datasets, with over 14000 genes having expression variation. **C.** Heatmap showing the top 50 differentially expressed genes between W2 and W110, including many histone genes and epigenetic regulators.

### Figure S3

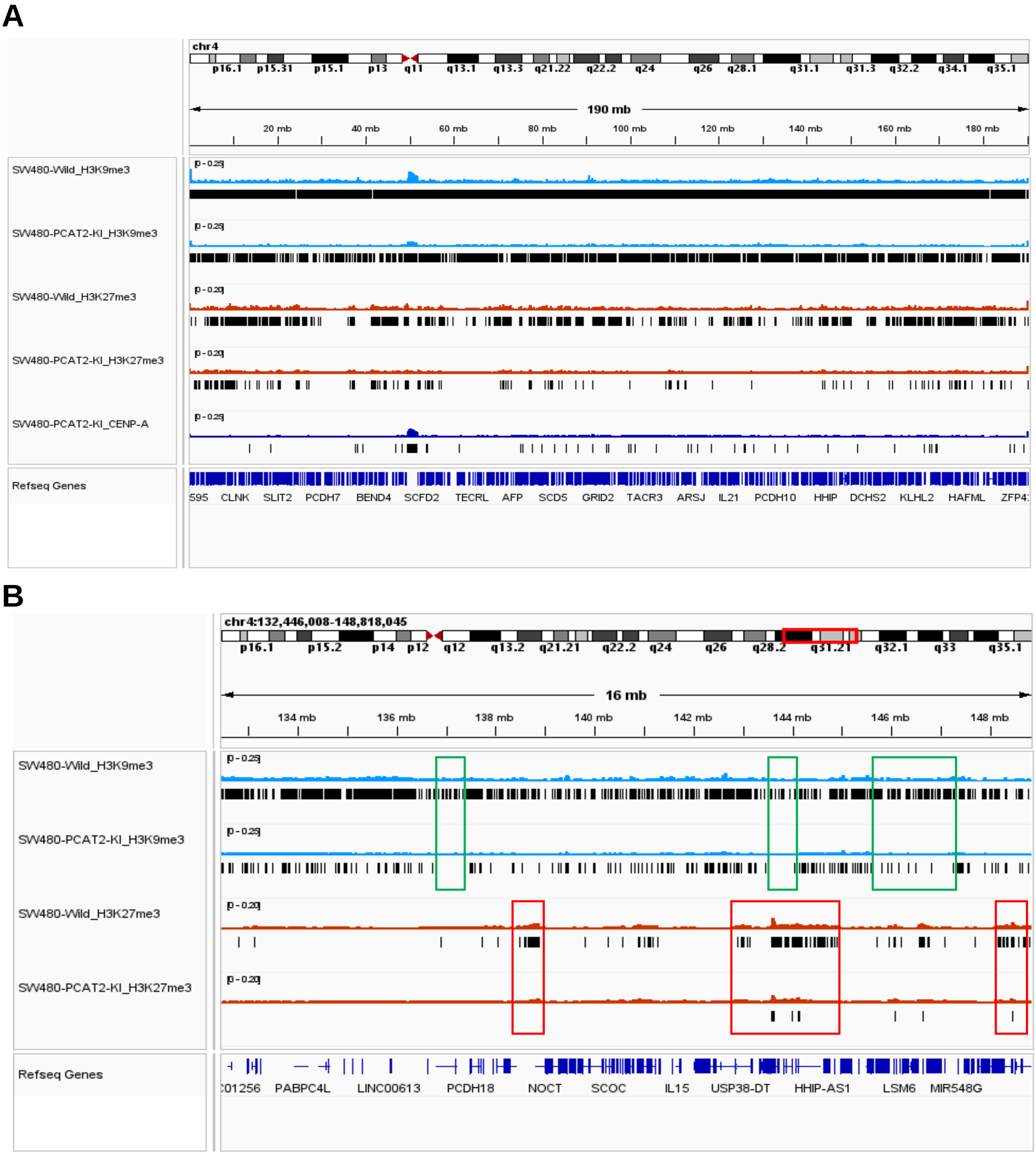


**Figure S3. Trans*PCAT2* gene insertion drastically altered epigenetic memory. A.** IGV browser screenshot of chromosome 4 of SW480$^{PCAT2\text{-KI}}$ showing significant loss of H3K9me3 and H3K27me3 through out the chromosome. The consensus peaks are marks as black bars below the corresponding ChIP-peaks. **B.** Chromosome 4q31 locus of SW480$^{PCAT2\text{-KI}}$ cells displaying a loss of H3K9me3 and H3K27me3 epigenetic signatures. With few H3K9me3 lost in the SW480$^{PCAT2\text{-KI}}$ cells' 4q31 locus compare to the SW480$^{\text{Wild}}$ cells (green box), a significant portion of the H3K27me3 signatures are lost in the SW480$^{PCAT2\text{-KI}}$ cells at this site (red box), specifically on up and down stream regions of the Trans*PCAT2* array knock-in site.

**Figure S4**

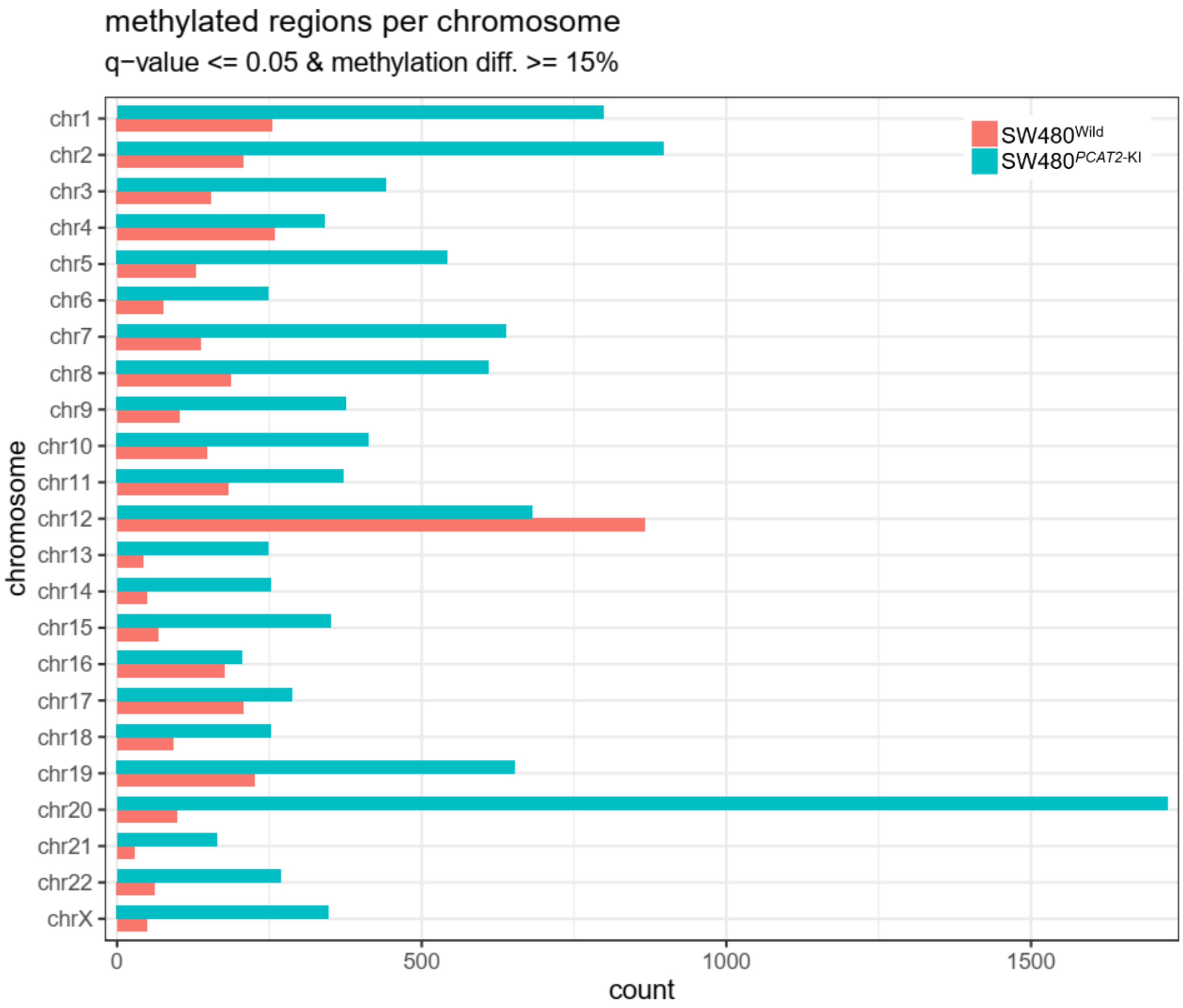


**Figure S4. Differential methylation signature of individual chromosomes.** Histogram showing the change in methylation profile of individual chromosomes in SW480$^{\textit{PCAT2}\text{-KI}}$ cells at W90 compared to SW480$^{\text{Wild}}$ cells.
