## Supplemental Table 1 for "Oncogenic lncRNA transgene transcription modulates epigenetic memory at a naïve chromosomal locus"

**Table S1.**

List of primer sequences used in this study.

| Gene | Experiment | Sequence (5'->3') |
|---|---|---|
| PCAT2 prmtr-1 Fwd | ChIP-qPCR | GGGACTTTCCTACTTGGCAGT |
| PCAT2 prmtr-1 Rev | ChIP-qPCR | TATCCACGCCCATTGATGTA |
| PCAT2 prmtr-2 Fwd | ChIP-qPCR | TACATCAATGGGCGTGGATA |
| PCAT2 prmtr-2 Rev | ChIP-qPCR | GTCAATGGGGTGGAGACTTG |
| MAML3 prmtr-1 Fwd | ChIP-qPCR | CGCTGTTACTAGCGCAGACTT |
| MAML3 prmtr-1 Rev | ChIP-qPCR | TGGAAAGAAAAAGGCAAATGA |
| MAML3 prmtr-2 Fwd | ChIP-qPCR | CACACTCACACACGCTCTCA |
| MAML3 prmtr-2 Rev | ChIP-qPCR | GCGACGGAGAGAAGAAAGATT |
| SCOC prmtr-1 Fwd | ChIP-qPCR | AAGGGCTAATGAAGCCAACA |
| SCOC prmtr-1 Rev | ChIP-qPCR | AACAGCAAGGTCTCCCTTTTC |
| SCOC prmtr-2 Fwd | ChIP-qPCR | TTCTTCCCCCAACCTTATCC |
| SCOC prmtr-2 Rev | ChIP-qPCR | ACCCTGCCTTCAAGTTTCCT |
| ZNF510 prmtr Fwd | ChIP-qPCR | TGGTGAAACCCCGTCTCTAC |
| ZNF510 prmtr Rev | ChIP-qPCR | CTCAGCCTCCCAAGTAGCTG |
| GAPDH prmtr Fwd | ChIP-qPCR | CAATTCCCCATCTCAGTCGT |
| GAPDH prmtr Rev | ChIP-qPCR | GCAGCAGGACACTAGGGAGT |
| Sat2 Fwd | ChIP-qPCR | CATCGAATGGAAATGAAAGGAGTC |
| Sat2 Rev | ChIP-qPCR | ACCATTGGATGATTGCAGTCAA |
| PCAT2 Fwd | RT-qPCR | CCCTTAAGGCACTGATGCTC |
| PCAT2 Rev | RT-qPCR | GTGCTGATGCCTCTGGAAAT |
| TransPCAT2 Fwd | RT-qPCR | TCGACTACTACGCTCGTCAGACTAC |
| TransPCAT2 Rev | RT-qPCR | GTGTTCCCTCCAAAATTCAGGTACT |
| cMYC Fwd | RT-qPCR | AATGAAAAGGCCCCCAAGGTAGTTATCC |
| cMYC Rev | RT-qPCR | GTCGTTTCCGCAACAAGTCCTCTTC |
| MAML3 Fwd | RT-qPCR | ATCAGCCCATGGCTTACGCT |
| MAML 3 Rev | RT-qPCR | GTGGTTCGGGGAAGTCCAAC |
| SCOC Fwd | RT-qPCR | TGCAACACTCAGGTCTGAAAA |
| SCOC Rev | RT-qPCR | AATGTGCTGTCTTCCTCCTCC |
| SCOC-AS1 Fwd | RT-qPCR | GCCCTGCTCAGAAGACACAA |
| SCOC-AS1 Rev | RT-qPCR | AGGGAGGGTCTGAGTAACCC |
| SMAD1 Fwd | RT-qPCR | TGAACCATGGATTTGAGACAG |
| SMAD 1 Rev | RT-qPCR | ACATCCTGGCGGTGGTATT |
| PLK4 Fwd | RT-qPCR | AGGGACTGCGTGAAGGAAG |
| PLK4 Rev | RT-qPCR | TTTCCAACTTTAAAATCCTCGATCT |
| DHX15 Fwd | RT-qPCR | GCCTCGTCGAAGTACTGACT |
| DHX15 Rev | RT-qPCR | CGTTCTAAATGTGCCACCTGC |
| GAPDH Fwd | RT-qPCR | GCGGTTCCGCACATCCCGGTAT |
| GAPDH Rev | RT-qPCR | CCCCACGTCGCAGCTTGCCTA |
