## Supplemental Table 2 for "Oncogenic lncRNA transgene transcription modulates epigenetic memory at a naïve chromosomal locus"

**Table S2:** *p*-values of all experiments.

Figure 1C. Trans*PCAT2* expression in SW480$^{PCAT2\text{-KI}}$ cells

| Week 16 vs. 90 | Week 90 vs. 110 | Week 16 vs. 110 |
|---|---|---|
| 0.0003 *** | 0.0012 ** | 0.0004 *** |

Figure 2B. Expression of 4q31 locus genes in SW480$^{PCAT2\text{-KI}}$ cells

| Gene | Week 1 vs. 90 | Week 1 vs. 110 |
|---|---|---|
| ***SMAD1*** | 0.5891 | 0.0002 *** |
| ***PLK4*** | 0.2576 | 0.0195 * |
| ***MAML3*** | 0.0005 *** | 0.0077 ** |
| ***SCOC*** | 0.0003 *** | 0.0293 * |
| ***SCOC-AS1*** | 0.0195 * | 0.0046 ** |
| ***DHX15*** | 0.0245 * | 0.0030 * |

Figure 2D. Fold enrichment of epigenetic marks

| Epigenetic marks | Fold enrichment in SW480$^{PCAT2\text{-KI}}$ cells | ***p-value*** |
|---|---|---|
| **H3K27ac** | 0.5688 | 0.0002 *** |
| **H3K27me3** | 1.420 | 0.0398 * |
| **H3K9me3** | 0.7831 | 0.0042 ** |

Figure 3E. Quantitative analysis of (ChIP-qPCR) H3K27me3 levels at target gene promoters

| Gene | H3K27me3 |
|---|---|
| **Trans*PCAT2*_pmtr-1** | 0.0001 **** |
| **Trans*PCAT2*_pmtr-2** | 0.0581 |
| ***MAML3*_pmtr-1** | 0.0593 |
| ***SCOC*_pmtr-1** | 0.0001 **** |

Figure 3F. Quantitative analysis of (ChIP-qPCR) H3K9me3 levels at target gene promoters

| Gene | H3K9me3 |
|---|---|
| ***MAML3*_pmtr-1** | 0.0064 ** |
| ***MAML3*_pmtr-2** | 0.0037 * |
| ***SCOC*_pmtr-1** | 0.0002 *** |
| ***SCOC*_pmtr-2** | 0.0013 ** |
| ***ZNF510*_pmtr** | 0.0001 **** |
| ***GAPDH*_pmtr** | 0.0031 ** |
| ***Sat2*** | 0.0011 ** |

Figure 5A. SW480$^{PCAT2\text{-KI}}$ cells with CENP-C colocalizing foci after Forskolin treatment

| Locus | DMSO vs. Forskolin |
|---|---|
| **8q24** | 0.0715 |
| **Trans*PCAT2*** | 0.2212 |

Figure 5C. LSD1 colocalizing foci on target locus

| Locus | SW480$^{Wild}$ vs. SW480$^{PCAT2\text{-}KI}$ |
|---|---|
| **8q24** | 0.2182 |
| **4q31/Trans*PCAT2*** | 0.0001 **** |

Figure 5D. Expression of target locus genes in SW480$^{PCAT2\text{-}KI}$ and SW480$^{Wild}$ cells after Dox treatment

| Gene | SW480$^{PCAT2\text{-}KI}$ | | |
|---|---|---|---|
| | **Dox treated** | **Outcome** | **Loci** |
| ***PCAT2*** | 0.0003 *** | Down | 8q24 |
| ***cMYC*** | 0.0003 *** | Down | 8q24 |
| ***TransPCAT2*** | 0.0029 ** | Up | 4q31 |
| ***MAML3*** | 0.1644 | No change | 4q31 |
| ***SCOC*** | 0.0437 * | Down | 4q31 |
| ***SCOC-AS1*** | 0.0019 ** | Up | 4q31 |

Figure 5E. SW480$^{PCAT2\text{-}KI}$ cells with CENP-C colocalizing foci after Dox treatment

| Locus | DMSO vs. Dox |
|---|---|
| **8q24** | 0.0906 |
| **Trans*PCAT2*** | 0.3944 |
